# Phenotypic and Genotypic Antiviral Resistance Testing of HSV-1 Causing Recurrent Cutaneous Lesions in a Patient with DOCK8 Deficiency

**DOI:** 10.1101/808303

**Authors:** Amanda M. Casto, Sean C. Stout, Rangaraj Selvarangan, Alexandra F. Freeman, Brandon D. Newell, Erin D. Stahl, Alexander L. Greninger, Dwight E. Yin

## Abstract

Antiviral resistance frequently complicates treatment of herpes simplex virus (HSV) infections in immunocompromised patients. Here we review the case of an adolescent boy with dedicator of cytokinesis 8 (DOCK8) deficiency, who experienced recurrent infections with resistant HSV-1. We used both phenotypic and genotypic methodologies to characterize the resistance profile of HSV-1 in the patient and conclude that genotypic testing outperformed phenotypic testing. We also present the first analysis of intrahost HSV-1 evolution in an immunocompromised patient. While HSV-1 can remain static in an immunocompetent individual for decades, the virus from this patient rapidly acquired genetic changes throughout its genome.

## Introduction

Up to 10% of herpes simplex virus (HSV) infections in immunocompromised patients are due to acyclovir-resistant viruses [1]. Resistant HSV infections are particularly common among allogenic hematopoietic stem cell transplant (HSCT) patients, in whom they cause nearly half of all HSV infections [1]. In immunocompetent patients, resistant HSV infections are less commonly encountered but are responsible for 6% of all HSV keratitis cases [2].

The clinical management of antiviral-resistant HSV infections is complicated by a number of factors. Treatment options are limited by the small number of effective antivirals available and the toxicities associated with therapies like foscarnet, cidofovir, and interferon. The development of resistance may be rapid, emerging with as little as two days of drug exposure [3]. Finally, HSV is the only virus for which resistance testing is performed phenotypically in clinical settings. These tests have long turnaround times and high inter-laboratory variability [4] and are commercially available only for acyclovir, ganciclovir, and foscarnet.

Here we describe the case of an adolescent boy with hyper-IgE syndrome due to dedicator of cytokinesis 8 (DOCK8) deficiency and recurrent HSV-1 infections to demonstrate the clinical challenges associated with the treatment of antiviral-resistant HSV infections. We compare the results of phenotypic and genotypic antiviral-resistance testing for HSV-1 from this patient and conclude that results of genotypic testing are more consistent with his previous antiviral exposures and his clinical response to certain antiviral agents. We also conduct the first study of HSV-1 evolution in an immunocompromised host by examining the genomes of longitudinal samples from the patient. We find that the patient’s HSV-1 rapidly acquired changes throughout its genome, challenging the paradigm that HSV-1 is a slowly evolving virus and suggesting that high rates of resistance are likely to be problematic for any antiviral used to treat HSV, particularly in immunocompromised patients.

## Methods

### Sample Collection and Phenotypic Resistance Testing

The patient’s parents provided written consent for sample collection from the patient and his brother and for the publication of de-identified medical information. This study was approved by the institutional review board of the University of Washington. Twelve HSV-1 samples were collected from the patient and one sample was collected from his younger brother (also affected by DOCK8 deficiency) during a four-year period. All samples were collected from clinically apparent, cutaneous lesions. Samples were grown in culture to confirm the presence of HSV-1. Seven samples from the patient were sent to a commercial laboratory for phenotypic resistance testing.

### Genome Sequencing and Analysis

We sequenced the genomes of seven HSV-1 samples from the patient and one sample from his brother using a probe-capture next-generation sequencing technique described previously [5]. Consensus genomes and alignment files were generated from raw sequencing reads using a publicly available computational pipeline [5].

Consensus genomes were aligned to an HSV-1 reference sequence (strain 17, JN555585.1) [6] with the multiple alignment using fast Fourier transform (MAFFT) algorithm [7]. Terminal repeats and intragenic regions were removed from the alignment. Single nucleotide variants (SNVs) were called relative to the reference using Geneious [8] with manual review. Minor variants present in at least 10% of reads were also called using Geneious [8] with manual review. A phylogenetic tree was generated using MrBayes [9] with the default settings.

### Data Sharing

Consensus genomes have been submitted to GenBank under accession numbers MN401201 – MN401208.

## Results

### Phenotypic resistance testing frequently indeterminate or inconsistent with previous antiviral exposures

The patient was diagnosed with DOCK8 deficiency by genetic testing as a toddler and was maintained on prophylactic acyclovir for years. HSCT was considered for the patient but was ultimately declined by his family. From age 11 to 15 years, the patient experienced multiple culture-confirmed HSV-1 infections at different cutaneous sites (Figure 1A, B). The first of these infections involved the right parietal scalp, ear, and face. A viral sample from the scalp was sent for phenotypic resistance testing. Results were indeterminate for acyclovir, but showed foscarnet resistance, although the patient had no previous known exposures to foscarnet. The patient initially improved on acyclovir, but the scalp lesion later worsened, developing into a large pink exudative plaque histologically consistent with *herpes vegetans* (Figure 1C). A second sample was sent for phenotypic testing and this result indicated acyclovir resistance and foscarnet sensitivity. Topical 3% cidofovir cream was added to the patient’s antiviral regimen and the lesions subsequently began to improve.

**Figure 1:**
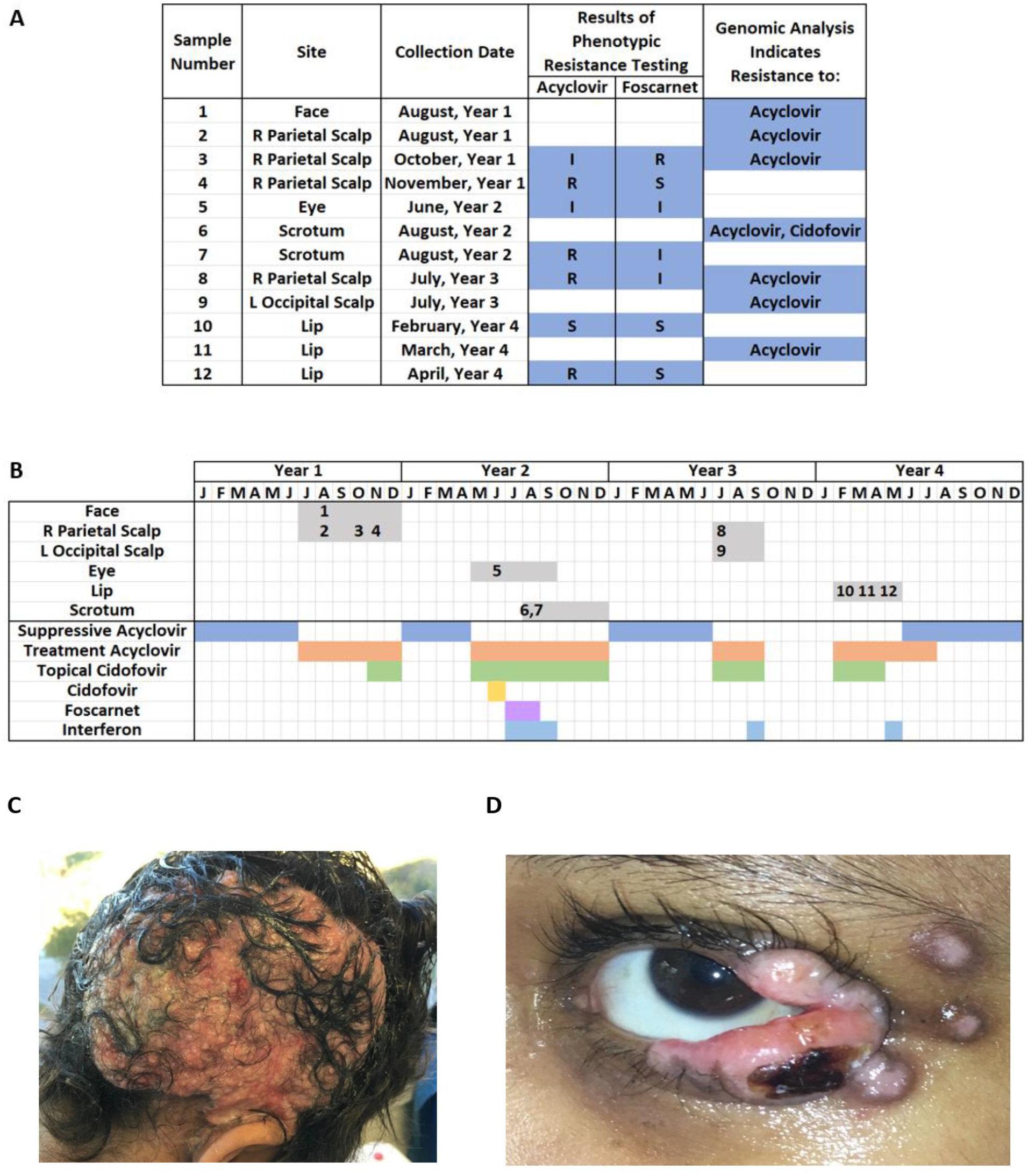
A) List of samples collected from the patient and the results of phenotypic and genotypic testing for resistance. For phenotypic testing, I = indeterminate, S = sensitive, and R = resistant. B) Timeline of HSV-1 infections and antiviral regimens. Numbers correspond to sample numbers in A. C) *Herpes vegetans* of right parietal scalp. D) Exophytic lesions adjacent to patient’s left eye.

Several months later, the patient developed purulent conjunctivitis of the left eye with adjacent exophytic papules (Figure 1D) and was treated with acyclovir. However, the periorbital lesions continued to worsen and systemic cidofovir was added. This regimen, too, failed to control the infection, so IV cidofovir was replaced with foscarnet and a dose of interferon alpha was administered. After these treatment changes, the eye lesions began to improve. Phenotypic testing of virus from the periorbital lesions was performed but resulted as indeterminate for both acyclovir and foscarnet.

As the eye lesions healed, the patient developed ulcerated nodules of the scrotum. Phenotypic testing of a sample from these ulcers showed acyclovir resistance but was indeterminate for foscarnet. The nodules were treated with a combination of acyclovir, interferon, topical cidofovir, and foscarnet. The latter two agents subsequently had to be stopped due to toxicity, but resolution of the nodules was eventually achieved with acyclovir and interferon.

For about six months after the scrotal lesions resolved, the patient had no evidence of active HSV-1 infection. The patient then developed a new lesion on the left occipital scalp and a smaller lesion on the right parietal scalp at the site of his previous infection. A sample from the right scalp was sent for phenotypic testing which indicated resistance to acyclovir but was indeterminate for foscarnet. The scalp lesions were treated with acyclovir and topical cidofovir with some improvement. However, the addition of interferon was again required to achieve resolution.

Between each of the four HSV-1 infections described above, the patient was maintained on low-dose, suppressive acyclovir. However, acyclovir was discontinued after resolution of the scalp lesions in hopes that the patient’s HSV-1 would become more susceptible. About five months later, the patient developed ulcerations and a pustule on his upper lip. Two samples were sent from these lesions for phenotypic testing. The first showed acyclovir sensitivity while the second demonstrated acyclovir resistance. Both resulted as sensitive to foscarnet. The lip lesions were initially treated with acyclovir and topical cidofovir. The latter subsequently had to be stopped due to toxicity and was replaced with interferon. After this substitution, the lip lesions began to improve.

### Genotypic testing for antiviral resistance consistent with history of antiviral exposures

To better understand the resistance profile of this patient’s HSV-1, we performed a retrospective genomic analysis on seven of the patient’s samples. Consistent with the patient’s chronic exposure to acyclovir, all sequenced samples were found to carry a single nucleotide consensus change in the thymidine kinase gene known to confer acyclovir resistance (c.527G>A, p.Arg176Gln) [10–12]. The only sequenced sample collected from the scrotal ulcers was also found to carry a single nucleotide consensus change in the DNA polymerase gene known to confer both acyclovir and cidofovir resistance (c.2462C>T, p.Thr821Met) [13]. This sample was collected one month after the patient was exposed to systemic cidofovir. No other genetic variants known to confer antiviral resistance were observed in any of the seven samples as consensus changes or minor variants [14].

### Samples from this patient rapidly accumulated genetic changes across the genome

We next examined the genic regions of the seven sequenced HSV-1 genomes in their entirety. We found that HSV-1 from the patient closely resembled HSV-1 from his younger brother. There were only 15 SNV differences between the brother’s sample and the first sequenced sample from the patient. The brother’s HSV-1 also carried the same acyclovir-resistance mutation in thymidine kinase that was observed in the patient. We noted that over time the patient’s HSV-1 samples became less similar to one another (Figure 2A) and to the brother’s HSV-1 sample, acquiring approximately four single nucleotide changes per year (Figure 2B). Anatomic site also appeared to play a role in the evolution of HSV-1 in the patient. The most divergent of the patient’s samples, both relative to the brother’s sample and to the patient’s other samples, came from the scrotal ulcers, which were anatomically isolated from all other lesions. This same sample was the only one from the patient to carry a cidofovir resistance mutation.

**Figure 2:**
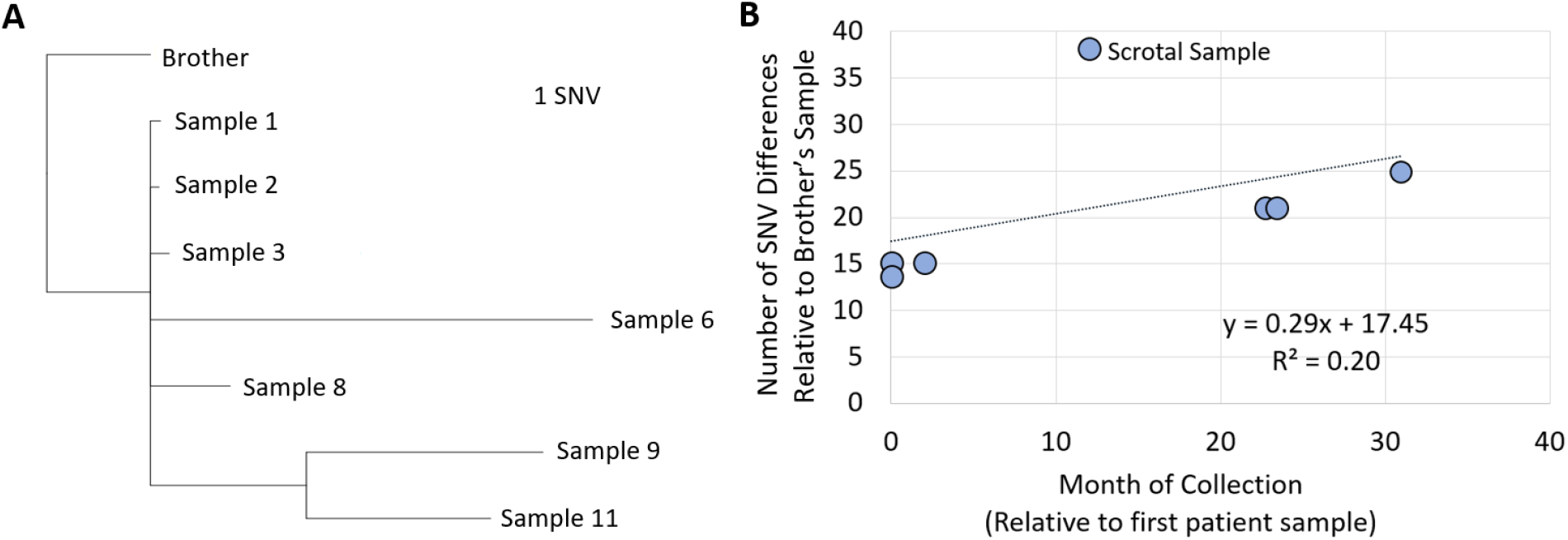
A) Phylogenetic tree of sequenced samples. B) Number of intragenic SNV changes relative to the brother’s sample graphed versus the time of collection for each of the seven sequenced samples from the patient.

## Discussion

Phenotypic testing is currently the only available method for assessing antiviral resistance in HSV in clinical settings. Its utility is limited by its long turnaround time (several weeks), its unavailability for some antivirals, and its frequent failure to provide definitive results. Four out of seven samples sent for phenotypic testing from the patient had indeterminate results for at least one antiviral and we were unable to test for phenotypic resistance to cidofovir, which was frequently used to treat the patient. Additionally, foscarnet resistance was reported for the first sample sent for testing. In retrospect, we suspect that this result was inaccurate, given that the patient had no previous exposure to this drug and that he subsequently responded well to it. Because of this result, we avoided using foscarnet to treat the *herpes vegetans* of the right parietal scalp and delayed its use for the periorbital lesions until other agents had been tried. Thus, inaccurate results from phenotypic testing can lead to avoidance or delay in administration of an effective therapy.

The main limitation of genetic antiviral resistance testing for HSV is that some variants in the thymidine kinase and DNA polymerase genes have not yet been phenotypically characterized. Nonetheless, in this case, genotypic testing predicted a resistance profile for the sequenced samples that was consistent with the patient’s previous exposures and responses to antiviral agents. In particular, the acyclovir resistance mutation found in all samples was consistent with the patient’s long history of exposure to the drug and explained why the patient’s infections did not respond well to acyclovir as monotherapy. Genotypic testing also indicated the presence of cidofovir resistance in the scrotal sample, consistent with the patient’s recent exposure to systemic cidofovir. Given these observations, we think that genotypic testing out-performed phenotypic testing for this patient, though we acknowledge that our assessment of clinical response to antivirals may have been confounded by the patient’s medical complexity and the effects of agents he received to treat other issues.

In addition to the information they provided about antiviral resistance, the HSV-1 genomes from the patient offered an unprecedented look at HSV-1 evolution in an immunocompromised host. HSV-1 can remain genomically static in immunocompetent hosts over decades [15]. This stasis stands in stark contrast to the multiple SNV changes that we observed in our patient’s samples. This difference in the rate of viral genomic change between immunocompetent and immunocompromised hosts is the likely genesis of the difference in the frequency of antiviral resistance between the two groups. The high mutation rate of HSV-1 in immunocompromised persons has important implications for the future management of antiviral resistance in HSV. First, it suggests that immunocompromised persons are likely to develop high rates of resistance to any antiviral, not just those that target thymidine kinase or DNA polymerase. Second, it suggests that immunocompromised hosts may not require prolonged exposure to an antiviral to develop resistance. Finally, we observed the presence of genetically distinct populations of HSV-1 at different anatomic sites in this patient and showed that such populations can have different antiviral resistance patterns, further complicating the management of HSV-1 infections in immunocompromised hosts.

In conclusion, we have demonstrated how the challenges of treating resistant HSV in immunocompromised hosts can be compounded by the limitations of phenotypic antiviral resistance testing. We have also demonstrated that genetic testing for resistance is a promising alternative, though one that has not yet been approved for clinical use. Finally, we have shown that in the context of immunocompromise, HSV-1 rapidly accumulates changes throughout its genome. These findings suggest that HSV may be capable of developing resistance to any antiviral in immunocompromised hosts, regardless of drug target, and explain the observed ability of HSV to quickly develop resistance in immunocompromised patients.

## Acknowledgements

We would like to thank this patient and his family for their courage, strength, and dignity in all that they endured.

## Meeting Presentations

This work was accepted for a poster presentation at the International Herpes Workshop held in Knoxville, TN in July 2019.

